# Estimation of fluid flow velocities in cortical brain tissue driven by the microvasculature

**DOI:** 10.1101/2024.12.23.630163

**Authors:** Timo Koch, Kent-André Mardal

## Abstract

We present a modelling framework for describing bulk fluid flow in brain tissue. Within this framework, using computational simulation, we estimate bulk flow velocities in the gray matter parenchyma due to static or slowly varying water potential gradients—hydrostatic pressure gradients and osmotic pressure gradients. Working with the situation that experimental evidence and some model parameter estimates, as we point out, are presently insufficient to estimate velocities precisely, we explore feasible parameter ranges resulting in a range of estimates. We consider the effect of realistic microvascular architecture (extracted from mouse cortical gray matter). Although the estimated velocities are small in magnitude (e.g. in comparison to blood flow velocities), the passive transport of solutes with the bulk fluid can be a relevant process when considering larger molecules transported over larger distances. We compare velocity magnitudes resulting from filtration and pulsations. Filtration can lead to continuous directed fluid flow in the parenchyma, while pulsation-driven flow is (at least partly) reversible. For the first time, we consider the effect of the vascular architecture on the velocity distribution in a tissue sample of ca. 1 mm^3^ cortical gray matter tissue. We conclude that both filtration and pulsations are potentially potent drivers for fluid flow.

**Summary (lay abstract):** Fluid transport through brain tissue is dynamic, but the basic properties of this flow and its variability are poorly characterised. Disturbed fluid transport was linked to Alzheimer’s disease. Moreover, fluid flow may be exploited to administer drugs to the brain. Using computer simulations, we estimate bulk flow velocities in the gray matter functional brain tissue due to changes in blood pressure and solute concentrations in unseen detail. We show that microvessel pulsations as filtration across the blood capillaries are potentially potent drivers of fluid flow in brain tissue.

## Introduction

Fluid transport through the brain parenchyma is dynamic, but fluid bulk flow velocity direction, magnitude, and variability are poorly characterized [12]. Irregular fluid transport is suggested to link to neurodegenerative diseases characterised by protein accumulation [14]. Moreover, the cerebrospinal fluid of the central nervous system is a potential pathway for drug administration to brain parenchyma as an alternative for administration via blood [31], circumventing the blood-brain barrier. The fluid-filled spaces of the central nervous system are shown schematically in Fig. 1. They include the large-scale fluid spaces in the spinal canal, around the brain, and in the ventricular system, as well as small-scale fluid spaces within the brain tissue.

**Figure 1:**
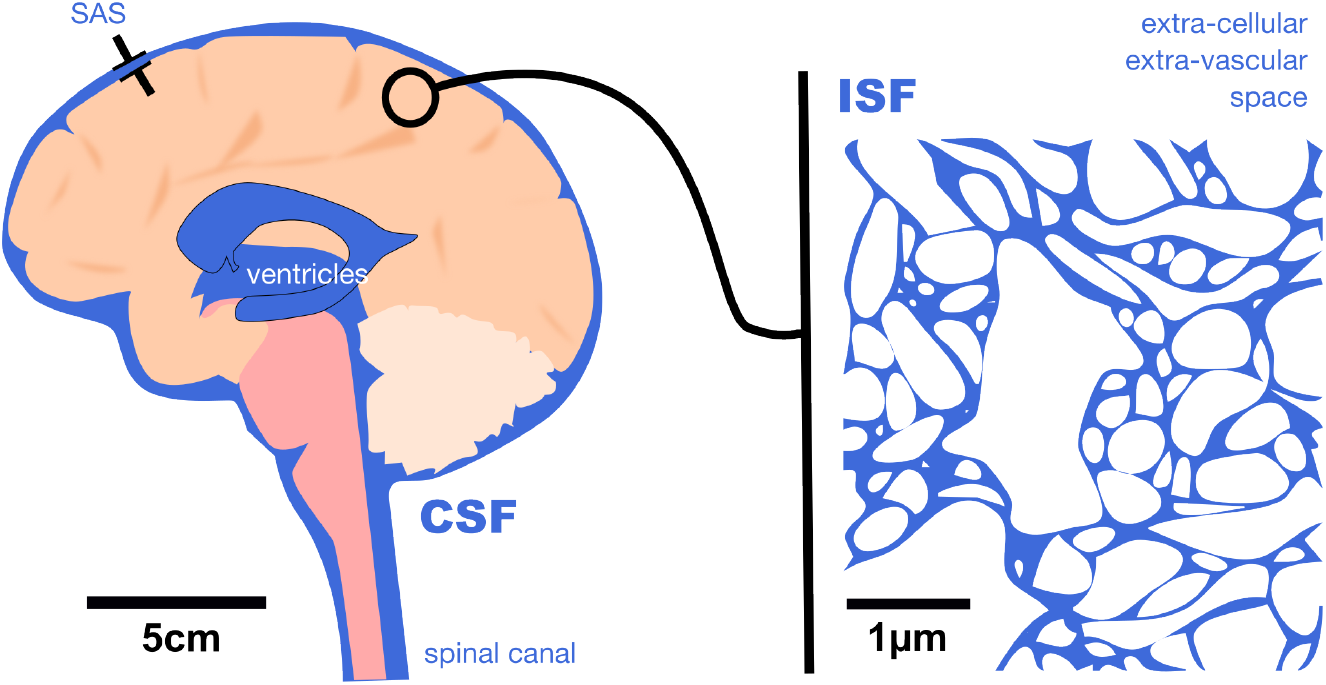
Fluid-filled spaces in and around the brain. Both cerebrospinal fluid (CSF, situated in the spinal canal, around the brain stem and in the subarachnoid space around the brain) and interstitial fluid (ISF, situated between cells in the parenchyma) are water-like fluids. CSF and ISF are close in composition and behave like water for the processes in the context of this article. The intracellular space (white on the right image) is also fluid-filled. The shown cartoon is only approximately to scale for a human brain, and the scale bars reflect the approximate spatial scale of the subfigures.

In this article, we attempt to estimate the potential of two processes to cause parenchymal fluid flow: water exchange between blood and extra-vascular tissue and microvascular pulsations. The former potentially causes directed continuous flow, while the latter may cause both oscillatory and directed continuous flow. In light of significant uncertainties regarding the magnitude of driving forces and their variability, we discuss a mathematical model of bulk flow that serves two purposes: stating significant parameters and boundary conditions that need to be measured and characterised better in the future, having a tool to study the influence of various processes and influence factors independently and in a controlled setting.

At the brain surface, experimental evidence in rodents demonstrates directed perivascular bulk flow inferred from observed micro-particle trajectories [28]. Similarly compelling evidence is lacking in the parenchyma. Studies with fluorescent or paramagnetic tracers remain inconclusive about the direction, magnitude, and variability of fluid bulk flow in the parenchyma [43, 10]. In such studies, the tracer is typically administered via the cerebrospinal fluid (CSF) or injected directly into the parenchyma and observed over time. Inferring bulk flow is complicated because tracer transport occurs due to several competing processes: passive transport with bulk flow (advection), molecular diffusion, and transfer to blood. Moreover, estimating concentrations from image intensity has its challenges, such as compartmentalisation of tracer, the influence of microstructural tissue heterogeneity, and, depending on the imaging technique, signal attenuation with cortical depth (imaging on µm scale), or effects of tracer dispersion by oscillatory velocity fields (imaging on mm scale).

To help enhance the understanding gained from indirect measurement techniques via tracer transport and in the lack of experimental techniques that directly quantify fluid transport on the relevant scale, computer simulations using mathematical models of bulk fluid flow built on physical principles make it possible to explore and test hypotheses for the underlying driving forces which are presently insufficiently characterised both in magnitude and spatial and temporal variability. Mathematical models make it possible to understand better the interplay and relative importance of various processes within the range of known parameters and boundary conditions. As many of the possibly relevant driving forces for bulk fluid flow are caused by interactions with the microvessels, we present in the following a model framework that allows to incorporate the discrete network architecture of microvessels to estimate flow velocities in the extra-vascular tissue between vessels.

Bulk fluid flow is both driven mechanically by hydrostatic pressure differences and by osmosis due to concentration differences across semipermeable membranes—the former results from blood vessels being pressurised in comparison with the surrounding tissue. Pulsatility of the heart (≈ 1 Hz in humans), lungs (≈ 0.3 Hz in humans), or active blood vessel dilation, i.e. vasomotion (≈ 0.1 Hz in humans and rodents) can additionally introduce local hydrostatic pressure gradients. Osmotic pressure gradients are induced by concentration differences in the fluid composition of cerebrospinal fluid, interstitial fluid, blood plasma, and intra-cellular fluid across semipermeable membranes such as the endothelial layer of microvessels. Concentration differences are caused by cell metabolism (such as neuronal activity) and are actively regulated by membrane channels. Moreover, bulk fluid flow in microchannels such as in the extra-cellular matrix can also be driven by electro-osmosis [42], an effect we will omit discussing in the present work.

The goal of this paper is threefold: (1) suggest mathematical models describing the basic physical processes on various scales; (2) estimate velocity magnitudes and discuss uncertainties; (3) point out parameters that introduce large uncertainty and where better quantification in terms of experimental data is needed. The developed mathematical models include and document the parameters and processes directly influencing bulk flow in brain tissue.

## Mathematical models for fluid flow

To the end of discussing influence factors and driving forces of flow, and conducting quantitative simulations, we present three models describing bulk fluid flow in brain tissue with different levels of accuracy, or in other words, on different spatial scales. We consider mathematical models on a microscale (nm to µm), a mesoscale (µm to mm), and a macro (organ) scale (mm to cm). Before introducing the models, we want to clarify that when discussing bulk flow velocity, different concepts are in use.

### Definitions of bulk flow velocity

When characterising and discussing fluid flow, it is essential to agree on a definition of the flow velocity. Since in the context of brain fluid transport, different velocity concepts are sometimes used interchangeably. We here try to emphasise the differences between notions of velocities depending on the spatial scale.

In general, velocity describes the speed of motion in a specified direction. Mathematically, the velocity is represented by a vector ***u*** ∈ ℝ^*d*^ (*d* is the space dimension, typically *d* = 3) with magnitude 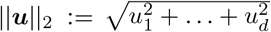 (the speed of motion) and direction ***u***||***u***||^−1^. Depending on the discussed spatial scale, we may distinguish several velocity definitions [2].

On the microscale (nm to µm), brain tissue is a heterogeneous medium with channel and tunnel-like structures (10 nm to 100 nm in diameter) of the extra-cellular matrix available for flow. We assume bulk fluid flow is amenable to a continuum mechanical description. The microscale velocity field, denoted by ***v*** in what follows, describes local variations in space and time in the fluid-conducting tissue compartments. Typical for viscous flow (a reasonable assumption for water-like fluids above a scale of a nm) is a locally parabolic velocity profile with largest velocity magnitudes in the middle of tunnels or sheets forming the extra-cellular space [19], cf. Fig. 2(right).

**Figure 2:**
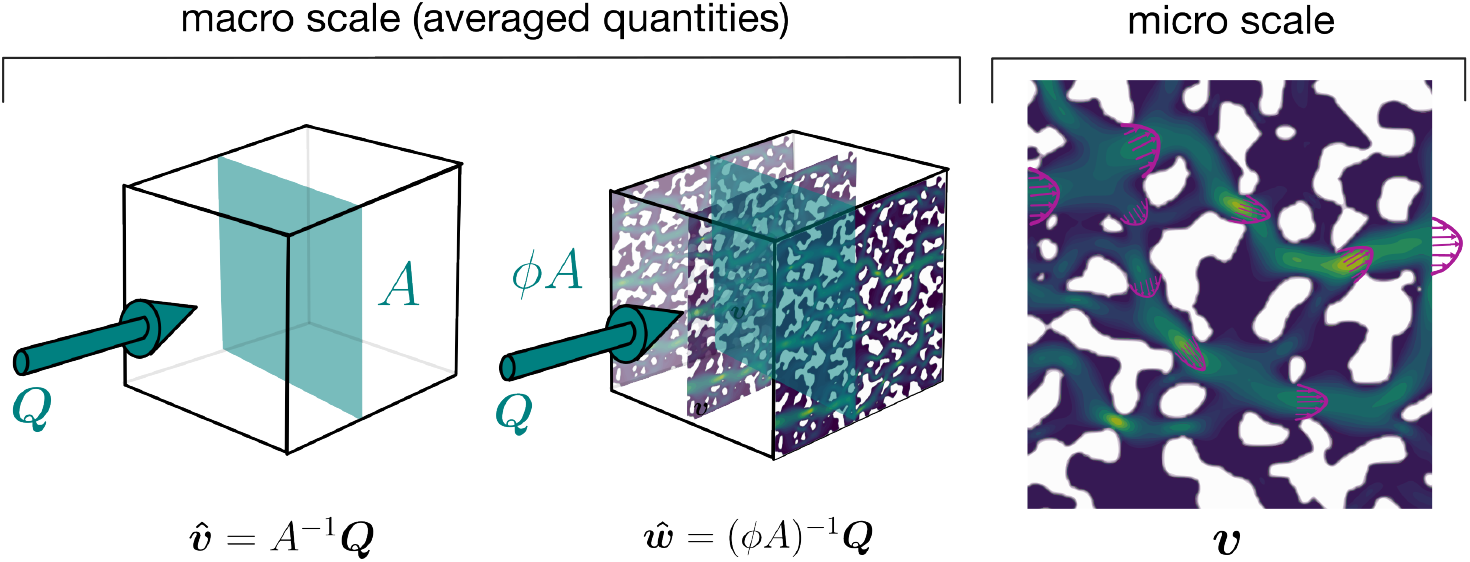
Different notions of bulk flow velocity. Right: The pore fluid microscale velocity field ***v*** describes the local velocity field between cells. Due to spatially varying pore space size, local velocity magnitude extrema can exceed or subceed the mean velocity by orders of magnitude. Left: On the macroscale, the macroscale velocity (superficial velocity) 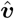 is implicitly defined as flow rate ***Q*** in a given direction through a surface with area *A*. Middle: The seepage velocity 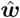 is a rescaled Darcy velocity through division by the flow-available volume fraction *ϕ* (porosity). It can used to estimate mean travel time.

On the mesoscale (*>* 3µm and *<* 1mm) and the macroscale (mm to cm), it is infeasible to resolve the local microstructure of extra-vascular tissue. Instead, we describe flow through the microstructure by volume-averaged quantities. Likewise, within the vasculature, it is typical to discuss cross-section averaged velocities. We denote averaged velocity fields by 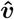 and say macroscale velocity, superficial velocity, or Darcy velocity. The velocity field 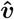 is to be interpreted as the area-specific flow rate in the direction of the unit normal vector on the chosen area element, cf. Fig. 2(left).

A third notion of velocity is given by the seepage velocity, or transport velocity 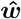. Since fluid only moves in the pore spaces, the average velocity in the pore space has to be higher than the area-specific flux. The transport velocity is estimated by 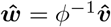, where *ϕ* is the volume fraction of the considered volume element (control volume) available for flow, cf. Fig. 2(middle). The transport velocity estimates travel time from one macroscopic point to another.

An example can illustrate the magnitude differences when discussing different velocities. Let’s assume a brain-scale (macroscale) model or macroscale experimental study (e.g. MRI) finds (by some method) an average macroscale velocity 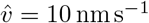 in cortical gray matter with direction towards the pial surface. This measure is useful, for example, to compute the flow rate over a given portion of the pial surface. However, interestingly, most of the bulk water does not move at this velocity. Since only the interstitial space is available for flow, the average transport velocity can be estimated by 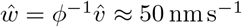, where *ϕ* = 0.2 (typical value for brain parenchyma [41]) is the fraction of space available for flow to total control volume. Due to variations in the local pore space geometry and alignment with the main flow direction, the local microscale velocity ***v*** in the interstitial space between two glial cells shows magnitudes up to two orders of magnitudes larger or smaller than the average such that we locally may observe 0 ≤||***v***|| ⪅ 5 µm s^−1^(typically following a skewed distribution with a long tail for large velocity magnitudes). Which velocity is relevant depends on the question asked.

### Microscale flow model (M_*µ*_)

Flow through a portion, Ω_*µ*_, of rigid^1^ interstitial space on the microscale is described by the incompressible Stokes equations

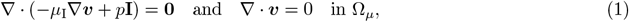

with the microscale fluid velocity ***v***, fluid pressure *p*, dynamic interstitial fluid viscosity *µ*_I_, the spatial gradient (∇) and divergence operators (∇·), and the identity matrix **I**. The momentum and mass balance equations, Eq. (1), are partial differential equations (PDEs) and are complemented by boundary conditions, either prescribing the velocity ***v***_*∂*Ω_, or the traction ***t***_*∂*Ω_, that is

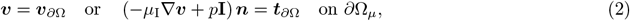

where *∂*Ω_*µ*_ denotes the fluid domain boundary. Equations (1) and (2) show that fluid flow can either be driven by the *deformation of the fluid domain boundary* (e.g. vessel wall deformation), a *nonzero boundary velocity* (e.g. filtration through the vessel wall), by a *normal stress* typically exerted by a pressure (e.g. blood pressure), or—less relevant in the current context—by a *shear stress*.

We omit here the description of a complete microscale blood flow model due to its complexity. However, we briefly discuss how to model filtration across the blood vessel wall. The endothelial cell layer forms the vessel wall separating the blood lumen and the perivascular space. The endothelial cell layer acts as a selective-permeable membrane. What distinguishes the endothelial layer of the microvasculature in the brain parenchyma from that of other organs is their virtual impermeability for ions [12]. This means ion transport is predominantly through ion channels that actively (e.g. by use of ATP to generate a cross-membrane electrochemical gradient) transport ions. In comparison, water can cross through endothelial tight junctions, diffuse through the cell membrane, or leak through membrane channels, and thus cross the endothelial cell layer easily in comparison.

As discussed in the extensive review by Hladky and Barrand [12], measuring local concentration in the vicinity of the vessel in vivo remains unattainable. However, comparing CSF and blood concentrations as shown in [12, Tab. 2] gives an estimate of the largest imaginable local differences (Δ*c ≈* 1.6 mmol kg^−1^ in humans). In the lack of precise data, we cannot quantitatively simulate water transport across the membrane. Instead, we here intend to give an upper estimate of the driving force and model the hypothetically resulting velocity field to get an idea of the magnitude and local variations in the parenchyma. To do so, with reasonable accuracy (in the sense of spatial resolution of the simulation results) and effort (in the sense of computation time), we next propose a model on a mesoscale.

### Mesoscale flow model (M_m_)

On the mesoscale (µm to mm), we assume that we cannot resolve individual cells and pore spaces of the extra-cellular matrix, and therefore use a locally homogenised (or volume-averaged) description. We resolve the geometry of individual blood vessels but describe the flow field within blood vessels only in terms of the mean flow velocity 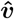. We denote the velocity magnitude with regular face 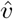 in the following. We choose a model description based on mixed-dimensional (1D-3D) PDEs, where blood flow is described on a network of segments, Λ, and interstitial flow is described in a three-dimensional domain, Ω, extended, also to include the space formally occupied by the vessels. The vessels and extra-vascular space coexist in this part of the domain, and the volume error made by this model assumption is assumed negligible due to a small blood volume fraction (*<* 5%). For a more detailed description of the model and a numerical solution strategy, we refer to [24, 21, 20] and Appendix C. In the general model description below, we also consider possible deformations of the vessel’s cross-sectional area. To this end, we assume that the deformations are small and slow in the sense that we have small Reynolds and Womersley numbers as typical for the microcirculation^2^. In the following, the subscripts *t* and *v* signify that a quantity belongs to the extra-vascular or vascular compartment, respectively. The subscripts *B* and *I* denote blood and interstitial fluid material parameters. The model describes flow using two coupled mass balance equations,

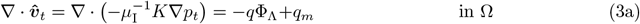

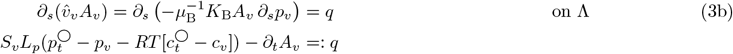

with extra-vascular macroscopic fluid velocity 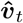 and average axial blood flow velocity 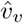, interstitial fluid and blood pressure *p*_*t*_ and *p*_*v*_ (in Pa). The coordinate *s* is the local axial coordinate in the vessel segment frame. The operator (*·)*^○^ denotes the average of (*·*) over the vessel perimeter with radius *r*_*v*_. The effective dynamic viscosity of interstitial fluid and blood are *µ*_I_ and *µ*_B_ (in Pa s). The effective blood viscosity, *µ*_B_, is modelled as a function of the local vessel lumen radius *r*_*v*_, see Appendix C and Table 1, whereas *µ*_I_ is assumed constant. Moreover, *K* and 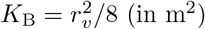 are permeability coefficients, the vessel perimeter is *S*_*v*_ = 2*πr*_*o*_, and *r*_*o*_ denotes the outer vessel diameter. The vessel lumen area is 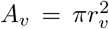. We take 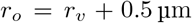. The filtration coefficient *L*_*p*_ (in m Pa^−1^ s^−1^) governs extravasation across an ideal semipermeable membrane, see Appendix A. The symbol Φ_Λ_ is to mean that the source term *q* in the three-dimensional PDE Eq. (3a) is distributed over the local surface of the vessel. (For more details on the derivation and implementation of the mixed-dimensional model chosen here, we refer to the literature discussing this method [24, 20].) The source term *q*_*m*_ is the metabolic water production rate in m^3^ s^−1^ m^−3^ (m^3^ fluid per second and per m^3^ tissue). Finally, 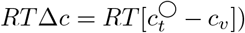 is the osmotic pressure difference between blood and ISF on the vessel surface (*R* is the universal gas constant and *T* temperature). Hence, hydrostatic pressure and osmotic pressure differences drive water exchange across the endothelial layer.

**Table 1:**
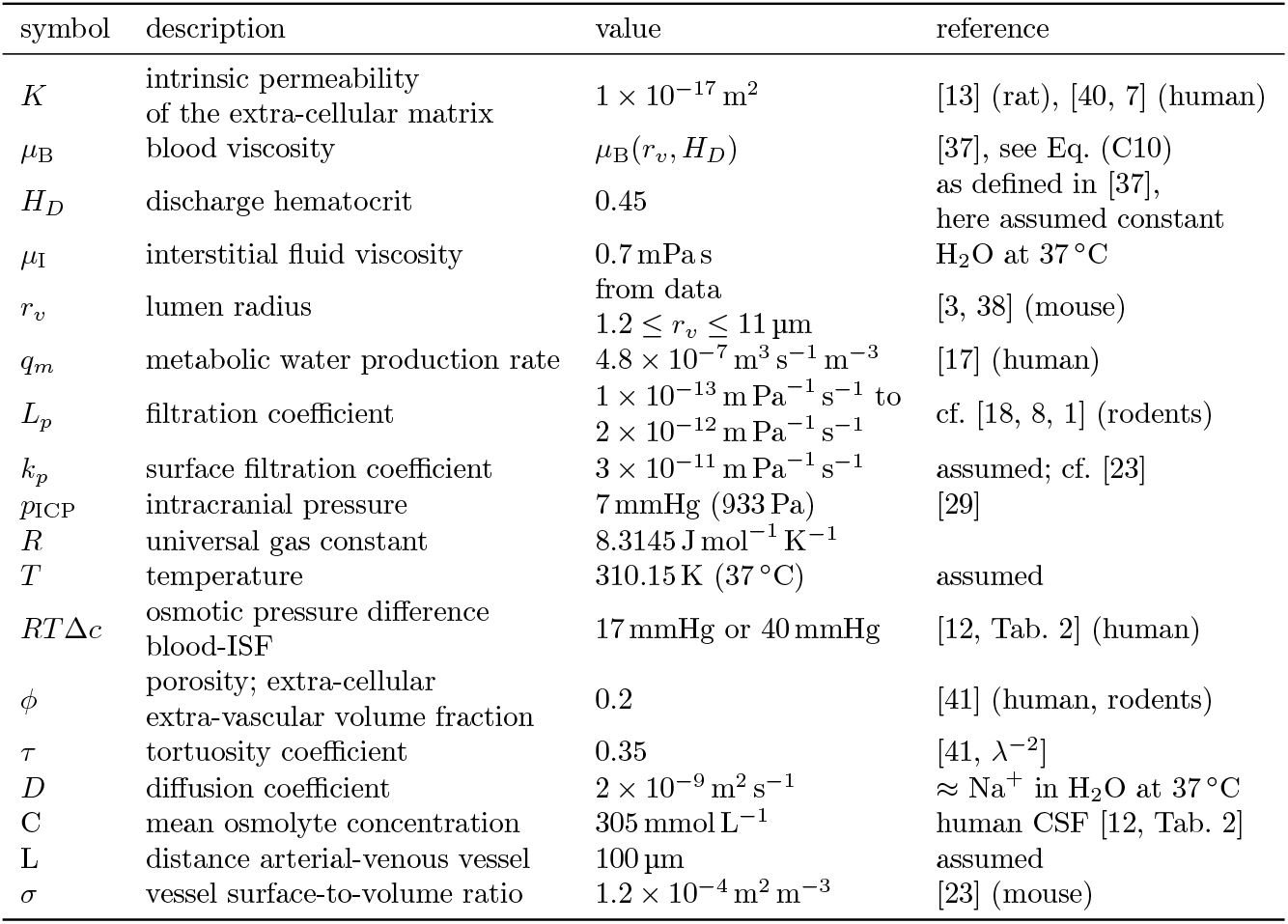
Parameters used in simulations with models M_*m*_, M_*mt*_.

Boundary conditions complement the balance equations. Let *∂*Ω denote the tissue domain boundary, and the points where the vessel centerline intersects the domain boundary are denoted by *∂*Λ. We set

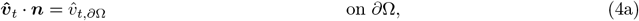

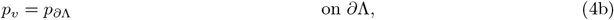

where 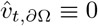 on all boundaries except for the pial surface where we model a porous membrane by prescribing 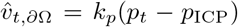, where *k*_*p*_ is a surface filtration coefficient and *p*_ICP_ is the intracranial pressure in the subarachnoid space above the tissue sample, and *p*_*t*_ is the extra-vascular fluid pressure evaluated on the pial-side boundary. The mesoscale model can describe local variations in blood flow velocity (i.e., from vessel segment to vessel segment) and interstitial flow velocity (i.e., velocity fields between vessels). However, it can not describe microscale variations (i.e., velocity fields between two individual cells).

### Mesoscale transport model (M_mt_)

To the end of estimating whether velocities predicted by M_m_ assuming a given osmotic pressure difference across the endothelial are large enough to cause an unstirred layer effect at the outside of the microvascular wall, we consider a second model that additionally models osmolyte transport using an advection-diffusion equation. Instead of using a mixed-dimensional approach, we only consider flow and transport in the extra-vascular space, Ω \ Θ, excluding the vascular space Θ, and model transport across the endothelial layer as a boundary condition. In the extra-vascular space, the equations governing bulk fluid and osmolyte transport for osmolyte concentration in the interstitial fluid *c*_*t*_ and fluid pressure *p*_*t*_ are given by

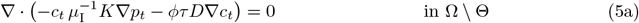

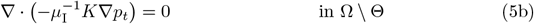

with boundary conditions such that the boundary is impermeable to osmolytes but permeable to water,

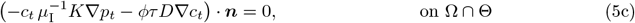

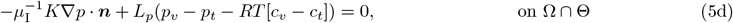

where *ϕ* is the extra-cellular volume fraction, *τ* the tortuosity coefficient, *D* is the binary diffusion coefficient of a given osmolyte in ISF, and ***n*** is a unit normal vector on Ω ∩ Θ pointing towards Θ. The osmolyte concentration and hydrostatic pressure in the blood, *c*_*v*_ and *p*_*v*_, respectively, are assumed to be given boundary values in this model. We point out that Eqs. (5b) and (5d) corresponds to Eq. (3a) with the source term in Eq. (3a) formulated as the boundary condition Eq. (5d).

### Macroscale flow model (M_c_)

By integrating the mesoscale model M_m_ over a larger portion of tissue—say a 1 mm^3^ cube—we can develop a two-compartment model that essentially states the integral fluid balance:

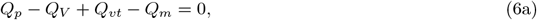

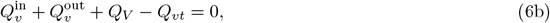

where 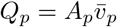 is the flow rate over the pial surface with pial surface area *A*_*p*_ and average Darcy fluid velocity 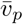, *Q*_*V*_ = *∫*_Λ_ *∂*_*t*_*A*_*v*_ d*s* is the flow rate due to changes in the vessel volume fraction *ϕ*_B_, *Q*_*m*_ is the endogenous water production due to cell metabolism,

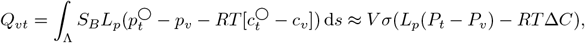

where *V* is the tissue sample volume, *σ* is the surface-to-volume ratio, *P*_*v*_ and *P*_*t*_ are the average vascular and extra-vascular fluid pressures, Δ*C* the average osmolyte concentration difference between the two compartments, 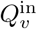 and 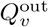 are the blood flow rates in and out of the given tissue volume. The model with the main symbols is depicted in Fig. 3.

**Figure 3:**
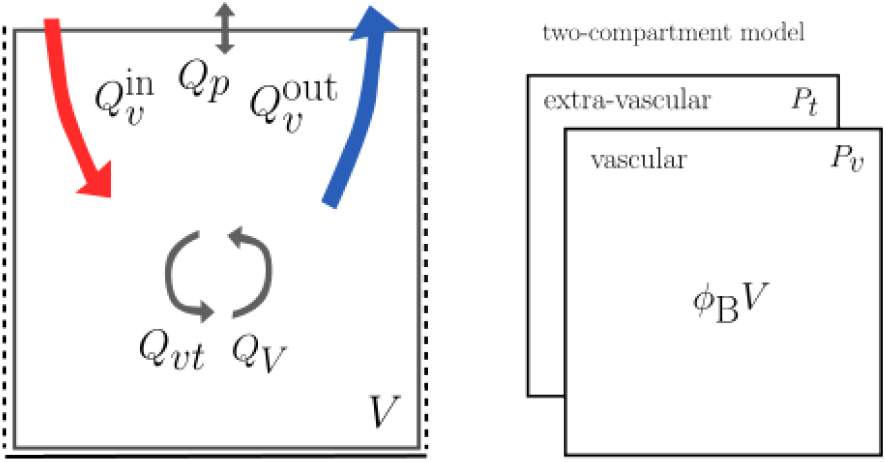
Macroscale flow model (M_c_). The macroscale flow model constitutes a global flux balance over a given portion of tissue conceptually split into two spatially coinciding compartments with given volume fractions.

Parameter values used in the simulations with all models are given in Table 1.

## Results

Holter et al. [13] used the microscale model (M_*µ*_), Eqs. (1) and (2), to estimate microscale interstitial flow velocities in a 64 µm^3^ cube of extra-cellular matrix (ECM)^3^ due to a prescribed pressure gradient of 1 mmHg mm^−14^ in one direction. They found a resulting mean microscale velocity magnitude of less than 10 nm s^−1^ with local velocities ranging from 0 nm s^−1^ to 50 nm s^−1^, cf. [13, Fig. 2]. The driving pressure gradient is assumed to be an upper estimate for typically occurring pressure gradients in-vivo without settling on a specific origin of the pressure gradient. This gives an idea of the order of magnitude of local velocities. However, two obvious shortcomings of such an analysis are (a) the driving forces have been postulated instead of motivated by a model, and (b) the direction of the driving force is geometrically simplified because the three-dimensional embedding of the microvascular network has not been considered.

### Velocities driven by the microvasculature

In the following, we discuss two possible driving forces for flow in the parenchyma: filtration via the endothelial layer or more generally the blood-brain barrier, and vessel dilation and restriction due to the heartbeat or vasomotion. In general, velocities that originate from either of these phenomena are higher near the vessel than at some distance from it. Following Darcy’s law, the magnitude of velocities caused by long and thin vessels acting as sources or sinks of fluid volume is proportional to the inverse of the radial distance from the vessel segment, ***v*** ∝*r*^−1^ (fluid pressure, *p* ∝ ln *r*) in the vicinity of the vessel. Therefore, velocities are expected to be higher near vessels than far from vessels. The actual velocity field results from an interaction of influences from all vessels surrounding an observed tissue location.

### Microvascular filtration may drive low magnitude directional flow

CSF is slightly hyperosmolar with respect to plasma: 1.6 mmol kg^−1^ [12] ∼ 5 ×10^3^ Pa. If this difference in osmolyte concentration persisted locally at the blood-brain barrier, it would drive water out of the blood into the interstitial fluid in addition to filtration due to hydrostatic pressure differences. Using the mesoscale model (M_m_) neglecting vessel pulsatility, we simulate the pressure and velocity field resulting from flow solely driven by imposing blood pressure values (spatially varying but constant in time) where vessels intersect the domain boundary. Appendix C describes the simulation setup in more detail. The model parameters are given in Table 1. We neglect pulsatility as we are interested in mean field filtration over longer time scales. Moreover, because of the endothelium’s low permeability, blood pressure fluctuations due to pulsatility cause negligible changes in filtration rates.

The resulting steady blood pressure field is shown in Fig. 4, blood pressures ranging from 5 to 56 mmHg. The cortical microvasculature forms a dense network. In the simulated microvascular domain, no point in the domain is further than 60 µm away from the closest vessel surface, cf. Fig. 4. This number is expected to be similar across species since oxygen’s characteristic diffusion length scale determines the inter-capillary distance. The resulting cerebral blood flow (CBF) measured as blood volume entering the domain via arterial vessels per minute and normalised by tissue volume is 95 mL*/*100mL*/*min. This value is the same for all simulation scenarios since the amount of water leaving the vasculature is less than 1 % of the total perfusion rate.

**Figure 4:**
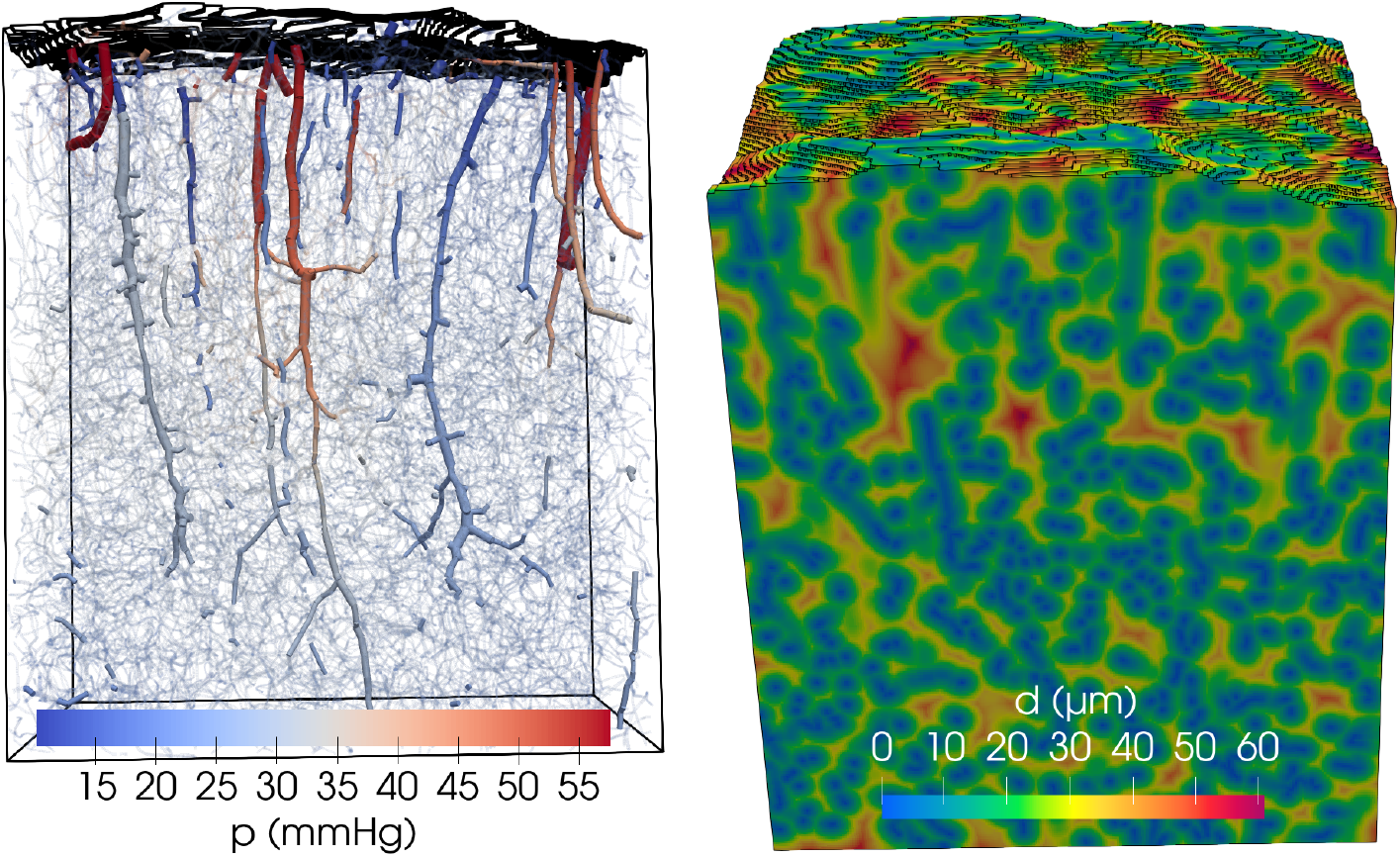
Pressure distribution in blood vessels (left) and tissue distance to the closest blood vessel (right). Pial vessels have been removed from the network, and the brain tissue domain follows the pial surface, estimated as a smooth envelope over the remaining vessels. Blood pressure boundary conditions are taken from a published data set [38], and the method for their estimation is described in [39]. The domain bounding box is 1 mm *×* 0.7 mm *×* 0.9 mm.

The velocity field for different osmotic pressure differences across the endothelial layer is shown in Fig. 5. The main trend in the velocity field is an upward direction with continuously increasing magnitude towards the pial surface due to the prescribed boundary conditions. Net filtration from blood and endogenous water production due to metabolism leads to a net source in the tissue. As the pial surface is set as the only permeable surface, the fluid has to move towards the pial surface where it exits into the subarachnoid space, which is kept at constant pressure *p*_ICP_ = 7 mmHg. Moreover, it can be observed that the larger the osmotic pressure difference the large the resulting velocities with maximum velocities at the pial surface of 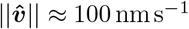 for *RT*Δ*c* = 40 mmHg (equivalent to a difference in osmolyte concentration of 1.6 mmol L^−1^), 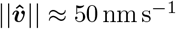 for *RT*Δ*c* = 20 mmHg (equivalent to a difference in osmolyte concentration of 1.6 mmol L^−1^), and 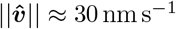 when assuming local osmotic equilibrium *RT*Δ*c* = 0. In the latter case, bulk fluid flow is driven by hydrostatic pressure differences and endogenous water production in the extra-vascular space. Assuming zero filtration and only endogenous water production of *q*_*m*_ = 4.8 *×* 10^−7^ m^3^ s^−1^ m^−3^ [17], the macroscopic flow model (M_*c*_) predicts *Q*_*p*_ = *Q*_*m*_. For a pial surface area of *A*_*p*_ = 1 mm^2^ and a tissue volume of *V* = 1 mm^3^, we get an average Darcy velocity of 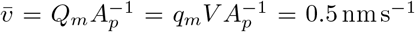. This is significantly lower than the velocity magnitudes estimated with filtration, and thus most of the fluid movement in Fig. 5C is predicted to be driven by hydrostatic extravasation.

**Figure 5:**
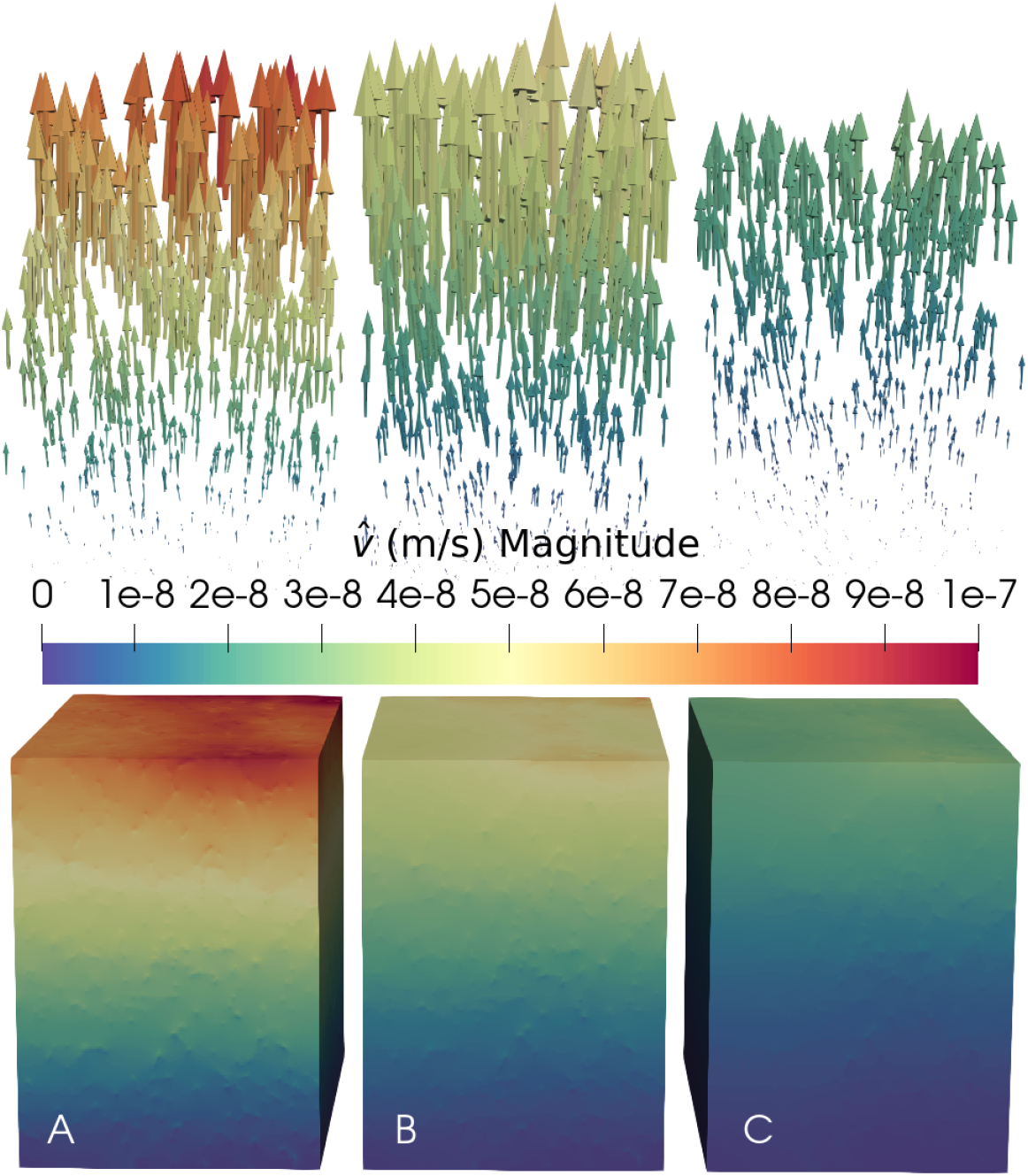
Simulation results with the mesoscale model M_m_ (varying osmotic pressure difference). Velocity field for *L*_*p*_ = 2 *×* 10^−12^ m Pa^−1^ s^−1^ and different osmotic pressure differences at the BBB: *RT*Δ*c*: 40 mmHg (≙1.6 mmol L^−1^) (A) 20 mmHg (≙0.8 mmol L^−1^) (B) 0 mmHg (C). The topmost tissue layer has been cut at 20 µm below the highest point of the pial surface for visualisation. (The pial surface is shown in Fig. 4.)

The velocity field for zero osmotic pressure difference and different filtration coefficients is shown in Fig. 6. We observe velocities at the pial surface of 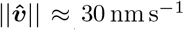 for *L*_*p*_ = 2 *×* 10^−12^ m Pa^−1^ s (same scenario as Fig. 5C), 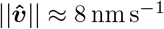 for *L*_*p*_ = 5 *×* 10^−13^ m Pa^−1^ s, and 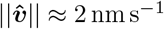 for *L*_*p*_ = 1 *×* 10^−13^ m Pa^−1^ s.

**Figure 6:**
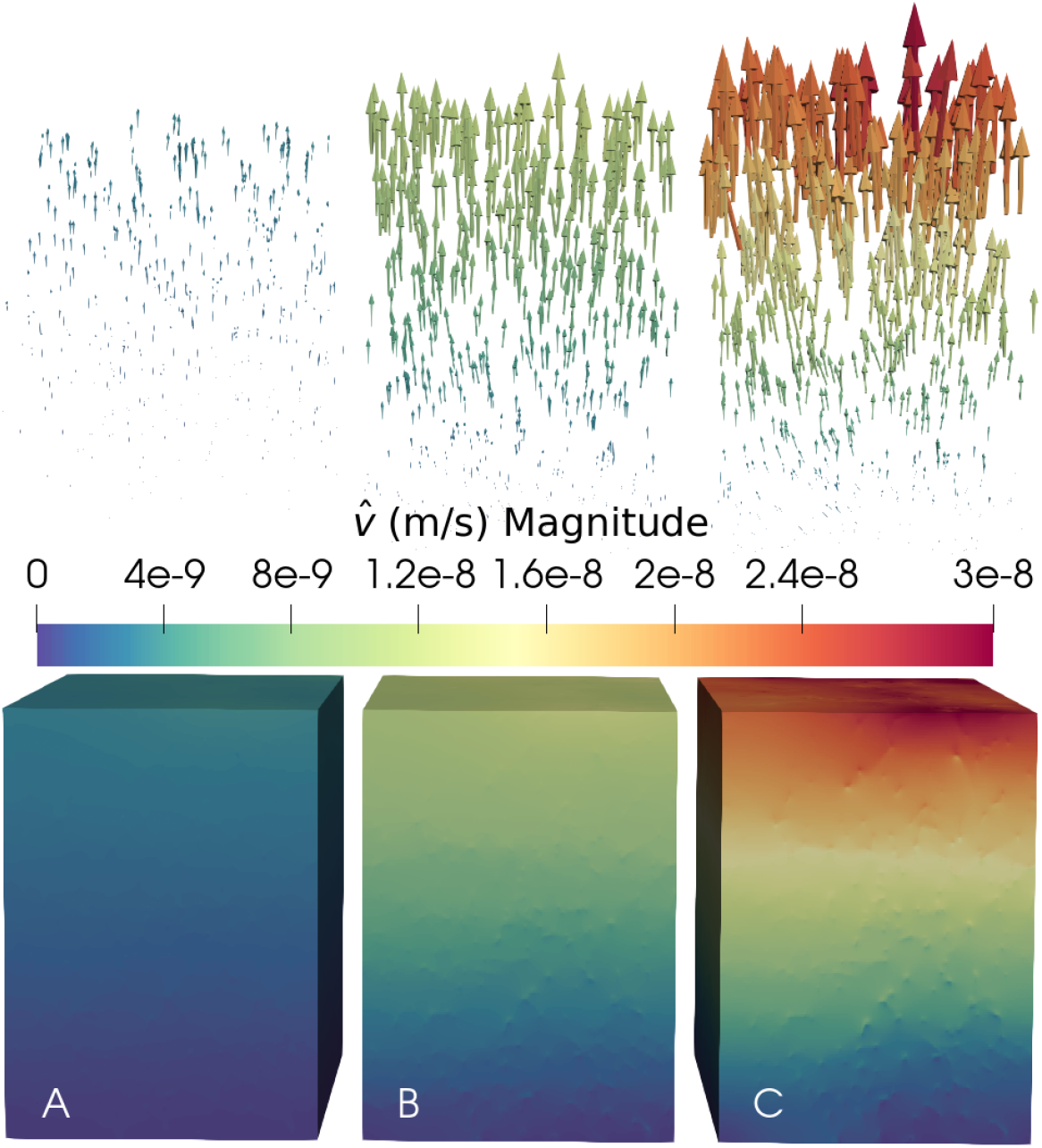
Simulation results with the mesoscale model M_m_ (varying filtration coefficient). Velocity field for zero osmotic pressure difference at the BBB and different filtration coefficients: *L*_*p*_ = 1 *×* 10^−13^ m Pa^−1^ s^−1^ (A) *L*_*p*_ = 5 *×* 10^−13^ m Pa^−1^ s^−1^ (B) *L*_*p*_ = 2 *×* 10^−12^ m Pa^−1^ s^−1^ (C). (C) is equivalent to Fig. 5C. The topmost tissue layer has been cut at 20 µm below the highest point of the pial surface for visualisation. (The pial surface is shown in Fig. 4.)

Figure 7 shows the horizontal velocity magnitude, omitting the vertical velocity component 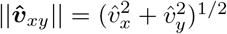, on horizontal slices parallel to the pial surface for different cortical depth. The presented result is obtained by assuming local osmotic equilibrium *RT*Δ*c* = 0 and *L*_*p*_ = 2 *×* 10^−12^ m Pa^−1^ s^−1^. Figure 7 demonstrates the local heterogeneity in the velocity field. The local velocity in the vicinity of vessels is an order of magnitude higher than the velocity in the space between. Each vessel perturbs the local flow field.

**Figure 7:**
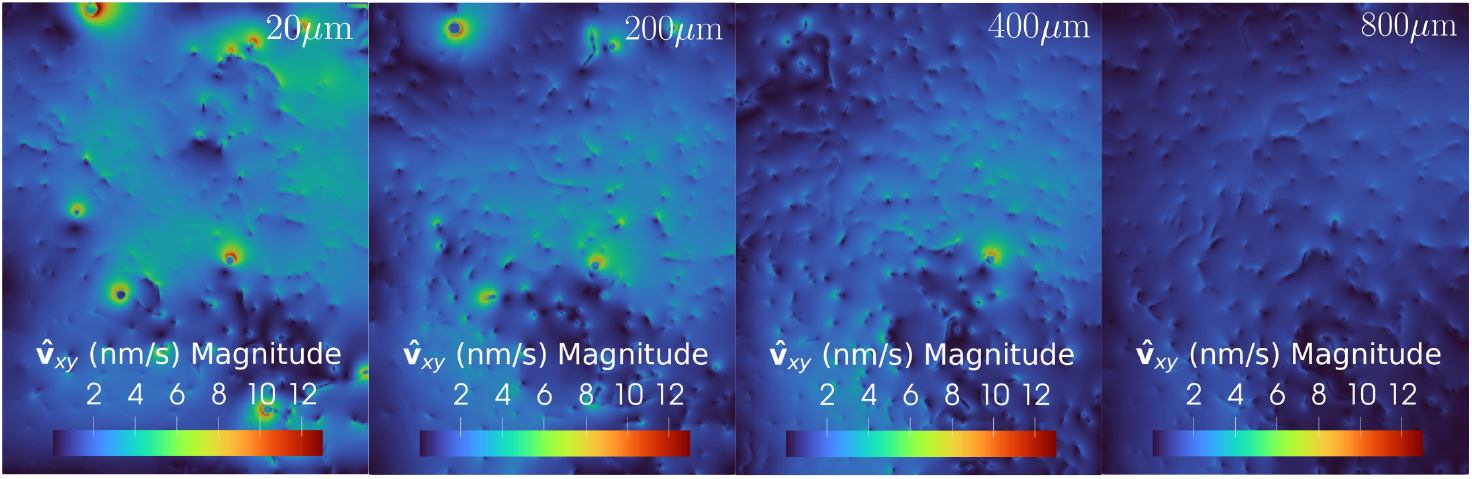
In-plane Darcy velocity magnitude as simulated with the mesoscale model. *M*_*m*_ shown on slices of different cortical depth (20 µm, 200 µm, 400 µm, 800 µm, where 0 µm is the pial surface). The velocity magnitude is computed based on the horizontal velocity components, neglecting the out-of-plane vertical velocity component. The slice dimensions are 0.7 mm *×* 0.9 mm.

### Unstirred layer effect reduces the potency of filtration as driving force

The unstirred layer effect describes an effect where downstream of the semipermeable membrane, the solvent motion drags along solutes such that the local concentration is lowered [32, 34, 12]. This effect reduces the effective osmotic pressure difference across the membrane and slows down transport across the membrane. The unstirred layer is expected to be stronger the higher the Péclet number, which relates the characteristic diffusive time scale to the characteristic advective time scale. The Péclet number is given by 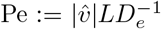 in which 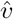 is the characteristic Darcy velocity magnitude, *L* a characteristic length scale, and *D*_*e*_ = *ϕτ D* the effective diffusion coefficient with extra-cellular volume fraction *ϕ* and tortuosity coefficient *τ*. With a small diffusion time scale (small Pe), diffusion can equilibrate local perturbations in the osmolyte concentration due to a source of liquid with a low concentration, as by filtration at the vessel wall.

Let us consider a simplified scenario as shown in Fig. 8 using the mesoscale transport model M_mt_. Assuming a distance between a major arterial microvessel and a major venous microvessel of *L* = 100 µm and small ionic solutes (*τϕD ≈* 1.4 *×* 10^−10^ m^2^*/*s (sodium ions in water at 37 ^°^C, *τ ≈* 0.34 [41], *ϕ ≈* 0.2 [41]), we can estimate Pe for different Darcy velocities. For the maximum velocity observed in Fig. 5, 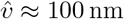, Pe = 0.07. Hence, diffusion is relatively important in comparison to advection. However, as we will demonstrate, the potency of the unstirred layer effect also depends on the absolute concentration of osmolytes present.

**Figure 8:**
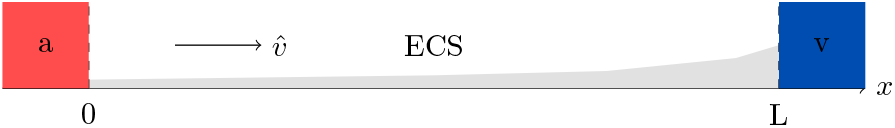
Simulation of the unstirred layer effect. Simplified setting for flow with Darcy velocity 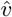 in the extracellular space (ECS) between two semipermeable membranes (endothelial layer, filtration coefficient *L*_*p*_) separated by parenchyma (ECS permeability *K*) of length *L* driven by the total pressure difference between an arterial location (a) and venous location (v). The velocity leads to a concentration gradient (shaded gray) against diffusion establishing a stable equilibrium situation with nonzero bulk fluid velocity. Large velocity would create a low concentration zone tissue-side at the inlet membrane, reducing the osmotic gradient across the membrane (unstirred layer effect). For small velocities, diffusion dominates advection and osmolytes in the ECS are well-mixed.

To discuss in more detail, we solve M_mt_ for the shown simplified one-dimensional scenario of Fig. 8. With Ω = [0, *L*] and spatially constant model parameters, see Fig. 8, Eq. (5) reduces to a nonlinear system of ordinary differential equations,

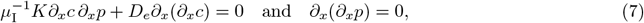

which admits the solution

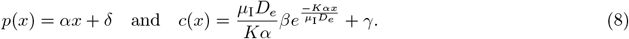

The constants *α, β, γ, δ*, are determined by no-flow boundary conditions for the concentration, conservation of osmolyte amount, and the filtration boundary conditions to blood at both ends, see Appendix B. The Darcy velocity is constant in this case, and in terms of the model parameters (see Appendix B), is given by

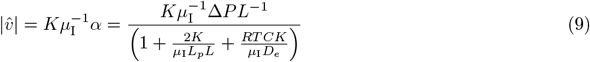

where Δ*P* is the total (osmotic and hydrostatic) pressure difference between arterial and venous vessels, *L* is the distance between the vessels, and *C* the average concentration, cf. Appendix B. The hypothetical velocity 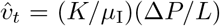 is the velocity that would result from the full total pressure gradient, Δ*P/L*, driving the flow in the interstitial space. As can be seen from equation Eq. (9), the actual velocity is reduced by two effects: the relative resistance of the vessel wall in comparison with the tissue permeability *K* (as quantified by the dimensionless term (2*K*)(*µ*_I_*L*_*p*_*L*)^−1^), and a reduction due to the unstirred layer effect (as quantified by the dimensionless term (*RTCK*)(*µ*_I_*D*_*e*_)^−1^). With parameter values in the range of Table 1, the term due to the wall permeability is ≈ 150, signifying that the vessel wall is a significant obstruction in line with the common assertion that the endothelial layer is virtually impermeable. Interestingly, the term due to the unstirred layer effect is ≈ 150, hence of equal order magnitude. In Fig. 9, we show the predicted relative reduction of the Darcy velocity due to the unstirred layer effect for different osmolyte concentrations (in the simplified setting above). Typical concentrations in parenchyma are on the order of 300 mmol L^−1^ [12, Tab. 2]. Hence, the unstirred layer effect may reduce the velocity to half its magnitude, and the velocity estimates given in the previous section are likely overestimations.

**Figure 9:**
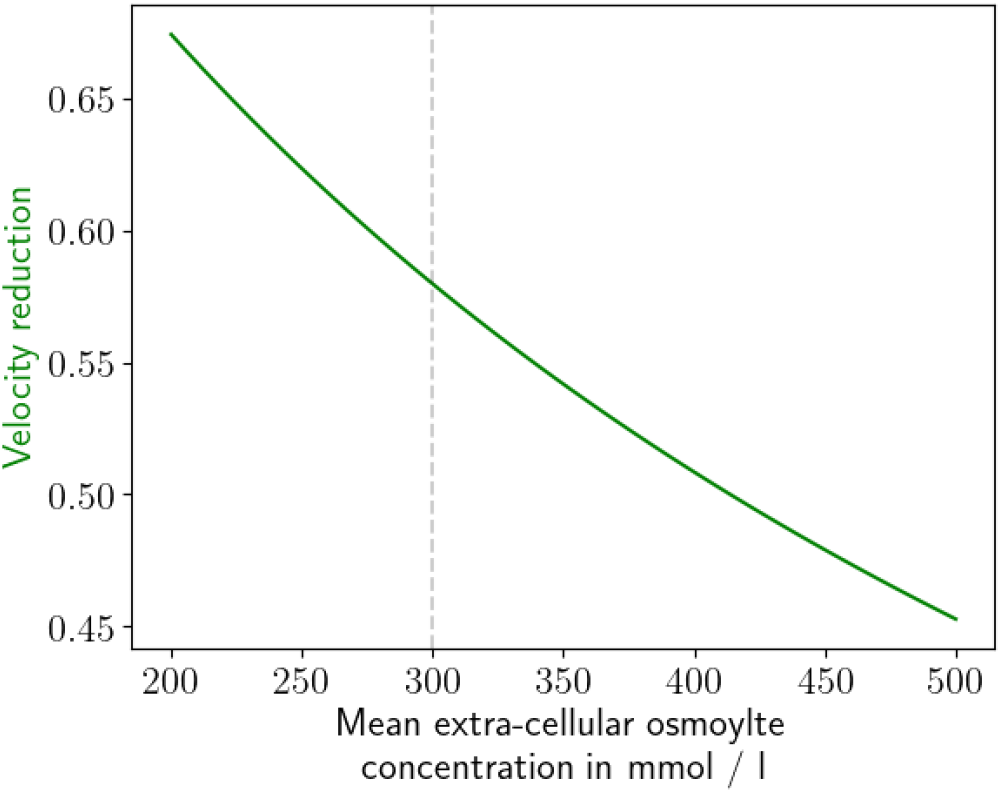
Unstirred layer effect. Reduction in velocity due to unstirred layer effect for different mean extra-cellular osmolyte concentrations.

### Pulsations potentially drive significant oscillatory flow

Microvessels are known to pulsate [27], both with the heartbeat (≈1 Hz) and during vasomotion caused by neuro-vascular coupling at lower frequencies (≈ 0.1 Hz) [4, 6]. Since brain tissue consists mainly of water and can therefore be considered incompressible, dilation of vessels will inevitably lead to fluid motion in the extra-vascular space.

To estimate the relative importance of pulsations and filtration on the local fluid velocities on the time scale of pulsation, we nondimensionalize with *ω*, the frequency of pulsations, *ζ*, the relative deformation amplitude such that the maximum outer radius *r*_0,max_ = (1 + *ζ*)*r*_*o*_, and Δ*P*, the characteristic total pressure difference across the membrane. We get a dimensionless number 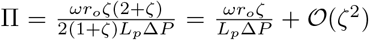 as the ratio of deformation velocity (*∂*_*t*_*r*_*o*_) to filtration velocities. For *ζ* = 0.01, *r*_*o*_ = 3 µm, *ω* = 1 Hz, *L*_*p*_ = 1 *×* 10^−13^ m Pa^−1^ s^−1^ to 1 *×* 10^−12^ m Pa^−1^ s^−1^, Δ*P* = 1 *×* 10^3^ Pa to 1 *×* 10^4^ Pa, Π = 3 to 300. This order of magnitude estimate shows that local velocities are likely to be significantly modulated by vessel pulsations.

We next consider a macroscale view (model M_c_ on a ≈ 1 mm^3^ with embedded microvascular networks as shown in Fig. 4. To get an idea of the magnitude of extra-cellular velocities that vessel pulsations may cause, we make the following assumptions leading to an *upper estimate* for the generated velocities: (a) fluid is only allowed to leave the domain through the pial surface, (b) all vessels dilate synchronously, (c) all components are incompressible.

In the absence of filtration and endogenous water production, the M_*c*_ extra-vascular balance equations give 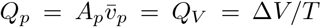, where Δ*V* is the total change of volume due to vessel dilation and *T* is the characteristic time scale over which this change occurs. For the considered microvascular network, cf. Fig. 4, a relative radius change of *ζ* = 0.01 leads to a volume change of Δ*V* = 0.2 nL, for *ζ* = 0.2, Δ*V* = 4 nL. Moreover, *A*_*p*_ = 0.63 mm^2^ (with *V* = 0.6 µL). The mean Darcy velocity at the pial surface can be computed as 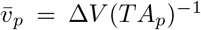. Consider *cardiac pulsations:* for a duration on the order of a systole, *T* ≈ 100 ms, synchronous cardiac vessels dilations (*ζ* ≈ 0.01) would cause an oscillatory velocity of amplitude 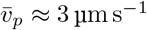. For comparison, compressing the entire gray matter tissue by 1 µm (≈ 0.1 % of the gray matter depth) within *T* = 100 ms yields 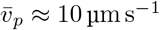. We remark that the cardiac motion is on the order of the resolution (≈ 1 µm for two-photon microscopy) of typical microscope measurements. So, even though these changes are small, they are hard to quantify even if high-resolution in-vivo imaging techniques are available, such as in rodents. Finally, we consider *vasomotion:* for a duration on the order of half a cycle, *T* ≈ 5 s [4, 6], vasomotion (*ζ* ≈ 0.2) would cause 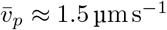 which is similar to cardiac pulsations.

## Discussion

Our computational analysis shows the potential of capillary filtration to act as a driving force for bulk fluid flow. Bulk fluid flow could, in turn, lead to enhanced clearance of tracers and metabolites out of the brain tissue. In a computational scenario, we assumed that an active regulation mechanism could uphold an osmotic pressure difference of up to 40 mmHg across the microvascular wall, with the extra-vascular space hyperosmolar to blood. We considered a cortical gray matter tissue sample of 0.6 mm^3^. The parameter choice leading to the largest filtration value is a larger osmotic pressure difference in combination with an upper estimate filtration coefficient *L*_*p*_ for a normal-appearing blood-brain barrier, cf. Fig. 5A. If such a voxel and the chosen parameter values were representative of the entire brain tissue of ca. 1200 mL, we obtain a resulting source of 9 L d^−1^. Gray matter amounts to about 65 % (ca. 800 mL) of brain tissue (55 % of total intracranial brain volume) in humans [26]. Assuming no filtration in the white matter tissue, the upper estimate would be reduced to 6.5 L d^−1^ which is still significant, and it seems unlikely that such a high rate would not have been detected, so far. Figure 5B (half the osmotic pressure difference) and Fig. 5C (no pressure difference) lead to 4.4 L d^−1^ and 2.5 L d^−1^ respectively when extrapolating our results for the cortical sample to the entire human brain. Since the model outcome is linear in the filtration rate, a filtration coefficient of *L*_*p*_ = 1 *×* 10^−13^ m Pa^−1^ s^−1^ (lower end of estimates) leads to 0.15 L d^−1^ and a coefficient of *L*_*p*_ = 5 *×* 10^−13^ m Pa^−1^ s^−1^ (mid-range) to a 0.7 L d^−1^ source when extrapolating our results for the cortical sample to the entire human brain. In these estimates, ca. 0.05 L d^−1^ [17] are attributed to metabolic water production. Water secretion in the choroid plexus is typically estimated to be ca. 0.5 L d^−1^ in humans. This comparison shows that filtration is a potentially potent driver of brain fluid movement. However, due to the high uncertainty in the model parameters, in particular the local osmotic pressure difference *RT*Δ*c*, and the filtration coefficient *L*_*p*_, quantitative conclusions are difficult and thus our estimates range from 0.15 L d^−1^ to 9 L d^−1^.

A compelling argument against filtration has been brought up, for example, by Hladky and Barrand [12]: filtration and the dilution of the extra-cellular fluid results in an adverse osmotic pressure gradient, which stops the flow. Our analysis of the unstirred layer effect suggests a possible way to circumvent this dilemma, given that extravasation velocities are small enough. While our results indicate that the unstirred layer effect likely reduces the filtration rate, the filtration rate is not reduced to zero. Given that there is a sink for fluid, which might be at some distant location to the microvessel surface or outside the brain tissue, a continuous bulk flow rate can persist in equilibrium with counteracting diffusion of osmolytes.

Since the suggested velocities are average Darcy velocities over the pial surface, local Darcy velocities at the surface may be much larger. Even larger velocity magnitudes (microscale velocity) could be attained locally within microscale fluid conduits. In our analysis, we have not considered that there may be highly conductive perivascular pathways, at least along the arterioles. In this case, much of the flow could be redirected to these channels, leading to relatively high local velocities [4]. Our obtained velocities are comparable to those reported for the perivascular spaces along pial vessels [28], making microvessel filtration one of the possible driving forces in consideration. Distinguishing quantitatively between perivascular and parenchymal flow would require the experimental characterisation of the geometric configuration of the perivascular space in the microvasculature in vivo, which is presently lacking.

The vascular architecture and several parameters in Table 1 stem from rodents and are not readily available or unknown for human tissue. However, we argue that they are expected to be relatively similar (considering the significant parameter uncertainties independent of species). The density of the capillary network is expected to be determined mainly by the characteristic length scale for oxygen diffusion, which is species-independent. Vessel diameters of similar order of magnitude, as is density with cortical depth where the average cortex depth is 2.5 mm in humans and 1.2 mm in mice [39]. Red blood cells are (on average) 55 fL in mice and 92 fL in humans, which affects the diameter of the smallest capillaries, which are larger in humans. Concerning pulsations, rodents have a much higher resting heart rate than humans, e.g. 10 Hz in mice and 1 Hz in humans. These interspecies differences add uncertainty to the estimates, and a more detailed investigation of interspecies differences would be necessary to quantify their effect on the estimated velocities. The presented methodology of estimation via mathematical models based on physical principles is unaffected by whether parameters correspond to humans or rodents and can, therefore, be used to quantify inter-species differences when available.

The direction of flow is prescribed by the boundary condition setup, and therefore, in our model, fluid leaving the simulation domain must leave towards the pial surface, leading to velocities directed towards the pial surface. At least circumstantial evidence [5] suggests that even tracer injected to the striatum in mice preferentially moved towards the pial surface. However, in experimental settings such as [5], one cannot easily distinguish between fluid-mediated transport due to filtration and pulsations or fluid motion caused by the tracer fluid injection. Better experimental measurements of spatial concentration maps during minutes after injection are required.

Based on the simplified macroscale model, the estimates regarding vessel pulsations show that vessel pulsations are a strong driving force for interstitial fluid flow. However, in the assumed model, flow due to pulsations is fully reversible and does not yield fluid net transport of fluids. For example, mechanisms of how symmetry may be broken have been proposed^5^ by Gan et al. [9]. Oscillatory velocity fields in tortuous porous media can lead to macroscale dispersion [33, 16] an effect that can be observed as enhanced Fickian diffusion or, depending on the local microstructural configuration and nature of interaction with the cells in the tissue also non-Fickian (anomalous) transport [25]. On the other hand, filtration leads to potentially reliable net transport given stable total pressure differences across the endothelial layer, albeit of smaller magnitude. In particular, for larger molecules and when considering transport over longer distances (e.g. 1 mm), even small bulk fluid flow can lead to enhanced transport.

A primary limitation of the presented analysis through computational models is that the required boundary data and parameters regarding local variations of pulsation and magnitude and variations of osmotic pressure gradients, in particular, are currently unknown ^6^. As such, no definite answer can be made about their relative importance. We consider the range of presented results informed guesses based on the currently available knowledge. Concerning the unstirred layer effect, we only investigated the mesoscale effect. However, a microscale view, cf. Fig. 10 on fluid transport, e.g. around endothelial tight junctions, could potentially reveal a more drastic picture of the unstirred layer effect if fluid flow is highly concentrated around such regions.

**Figure 10:**
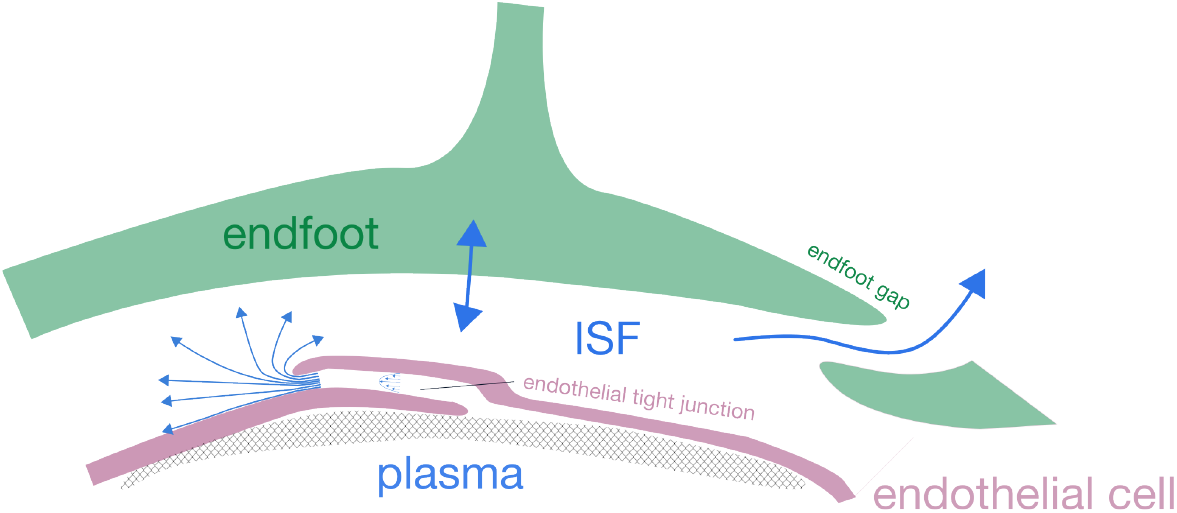
Simplified schematic of paravascular fluid exchange. (not to scale). Can local velocities cause an unstirred layer [32, 34, 11] such that the filtration rate is limited by diffusion? If yes, how large do local velocities have to be so this has a significant effect? Or how does the local microstructure affect the unstirred layer effect?

Finally, we want to remark that we have not considered ion movement models; indeed, the brain is a highly osmotically active environment with frequent local and global changes due to neuronal activity on various temporal and spatial scales. For instance, bulk flow due to local redistribution of charged ions in the inter- and intra-cellular space has been investigated with a mesoscale model in [42].

## Conclusion

Pulsations can potentially cause large velocities in comparison with blood filtration. The induced motion is likely oscillatory rather than continuously directional. Regarding tracer transport, filtration can cause advective transport, which is likely directed from the parenchyma towards the pial surface. Depending on the osmotic environment, which significantly influences velocity magnitudes but is not well characterised, such velocities may be relevant for large molecules when one is interested in transport over large distances. In combination with molecular diffusion, oscillatory flow driven by vessel pulsations can lead to dispersion on the macroscale, which appears as enhanced transport compared to pure diffusion in the direction of a negative concentration gradient. The presented mathematical model outlines the most significant unknown influence factors: spatial and temporal variability of pulsations, magnitude and spatial and temporal variability of osmotic driving forces, and the microvascular filtration coefficient. The currently available experimental evidence is insufficient to derive model parameters with the precision needed to make definite conclusions about the relative importance of bulk flow-generating driving forces. However, both pulsations and filtration appear to be potent drivers of interstitial bulk flow that shall be investigated with various methods.

## Acknowledgements

T.K. acknowledges funding from the European Union’s Horizon 2020 Research and Innovation Programme under the Marie Sklodowska-Curie Actions Grant agreement No 801133. T.K. and K.A.M. acknowledge funding by the Research Council of Norway, project 301013 (Alzheimer’s physics). K.A.M. acknowledges funding from the Stiftelsen Kristian Gerhard Jebsen via the K. G. Jebsen Centre for Brain Fluid Research and the European Research Council under grant 101141807 (aCleanBrain) and the national infrastructure for computational science in Norway, Sigma2, via grant NN9279K.

## Data availability

The network and boundary condition data is available at [38]. The software is available from https://git.iws.uni-stuttgart.de/dumux-pub/Koch2024a or via Software Heritage permalink https://archive.softwareheritage.org/swh:1:rev:393119cdcc33e049c82fadc3474b78de3af3ca3d.

## A Transport across semipermeable membranes

We follow Kedem and Katchalsky [15]. We denote temperature and pressure by *T* and *p*, and the ideal gas constant is *R* = 8.314 J K^−1^ mol^−1^ = 8.314 Pa m^3^ K^−1^ mol^−1^. The chemical potential difference (in J mol^−1^) between an inside (*i*) and outside (*o*) compartment separated by a membrane is modeled as

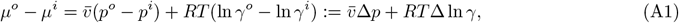

where 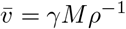 (in m^3^*/*mol) is the partial molar volume and *γ* the molar fraction of the constituent. Entropy production in a thermodynamic process has to be larger than or equal to zero. We define the dissipation function

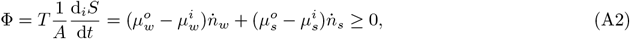

where *w* denotes the solvent, *s* the solute, *S* the entropy, and 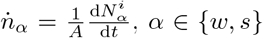 are flow rates per unit area across the membrane. For dilute binary solutions, ln *γ*_*s*_ ≈ *γ*_*s*_, ln *γ*_*w*_ = ln (1 − *γ*_*s*_) ≈− *γ*_*s*_. Combining the above equations, we obtain

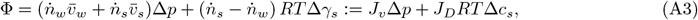

where *c*_*s*_ = *γ*_*s*_*V*_*m*_ is the solute concentration if *V*_*m*_ denotes the molar volume (in m^3^*/*mol) of the solution, and *J*_*v*_ and *J*_*D*_ are total volume flow rate and relative volume flow rate of solute and solvent. According to the theory of irreversible thermodynamics, *J* are functions of all driving forces (here Δ*p* and *RT*Δ*c*_*s*_) and linearly related, assuming the forces are sufficiently small,

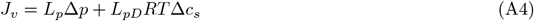

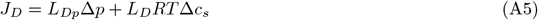

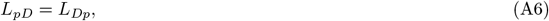

where the last equality is due to Onsager [30].

In the special case of an ideal semi-permeable membrane, 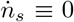, and thus 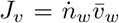 and 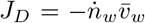for dilute solutions, hence *J*_*v*_ = −*J*_*D*_. Combining the equations for *J*_*v*_ and *J*_*D*_ above, we obtain the constraint

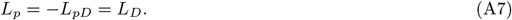

Hence, the system is entirely determined by a single filtration coefficient *L*_*p*_.

## B 1D mesoscopic osmolyte transport model

The nonlinear system of ODEs in 1D (Ω \ Θ = [0, *L*]) with (*·*)^*′*^ = *∂*_*x*_(*·*), *x ∈* [0, *L*],

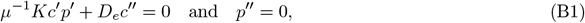

admits the solution

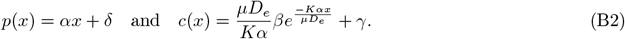

The constants *α, β, γ, δ*, are determined by no-flow boundary conditions for the concentration, conservation of osmolyte amount, and the filtration boundary conditions to blood at both ends:

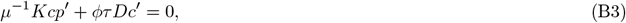

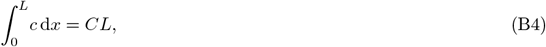

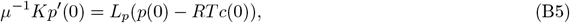

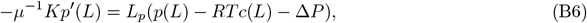

with the average osmolyte concentration *C*, the total pressure difference 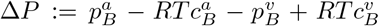 between arterial (*a*) and venous (*v*) blood, yielding *γ* = 0 and

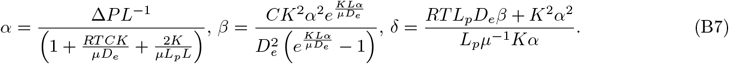

Note that the Darcy velocity magnitude is given by 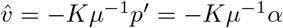.

## C 1D-3D mesoscale flow simulations

*×*

The blood pressure values are taken from the dataset [38], which constitutes a network with three-dimension embedding, vessel radii *r*_*v*_ for all segments, and blood pressure estimated in [39] by embedding the present network into a more extensive network. The microvascular networks approximately 7 *×* 10^5^ vessel segments have been extracted from the mouse brain cortex by [3]. The grid resolution is chosen so that no cells in the network are larger ≈ 3 µm by locally refining the mesh. The three-dimensional grid is Cartesian, with cells in the subarachnoid space removed and cell diameters of ≈ 3 µm. Blood flow in the microvascular networks follows Hagen–Poiseuille flow. Flow in the vascular network, and the extra-vascular tissue is solved in a fully coupled fashion as a single mixed-dimensional PDE problem using a method presented in [24, 21] and further numerically tested for the perfusion problem at hand in [20]. The technique is based on a mixed-dimensional finite volume solver implemented in the numerical software framework DuMu^x^ [22] (see “Code availability” for a link to the source code). The total number of degrees of freedom for the M_m_ simulation was 17 million. The resulting linear system after discretisation is solved with a block-diagonal algebraic multigrid preconditioned bi-conjugate gradient solver. The pressure boundary conditions were estimated in [39, 38]. Apparent blood viscosity was estimated by the non-linear in vitro apparent viscosity relation proposed by [37]. The empirical relationship was obtained for human blood samples where the mean red blood cell volume is 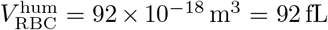. According to Pries and Secomb [35], a relation for other species with different red blood cell volumes may be obtained by multiplying the vessel diameters in the equation by a factor 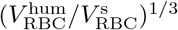. For mouse blood with 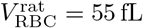, this gives a factor of 1.187 [36] which we used to match the lumen diameters in the employed mouse network geometry. Setting *D*:= 2 *×* 1.187*r*_*v*_ (where we recall that *r*_*v*_ denotes the local lumen radius), we assume *µ*_*B*_ = *µ*_*B*_(*D*(*r*_*v*_), *H*_*D*_) [37]:

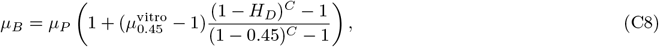

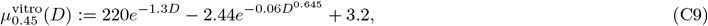

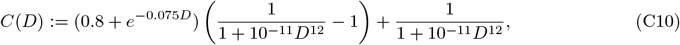

where *η*_0.45_ is the relative apparent viscosity at a discharge hematocrit of *H*_*D*_ = 0.45, *C* is a scaling function, and *µ*_*P*_ = 1.3 mPa s.

## Footnotes

1 in the sense of sufficiently small and slowly varying deformations of the fluid domain.

2 The Reynolds number for blood flow is given by 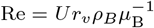 with a characteristic velocity *U* and the blood density *ρ*_*B*_. The Womersley number is given by 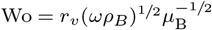 with the characteristic angular frequency *ω* (e.g. *ω* ≈ 1 Hz for a human heart beat). In the microcirculation, typically Re ≪ 1 and Wo ≪ 1.

3 The ECM geometry was reconstructed from electron microscope images of rat hippocampal neuropil by Kinney et al. [19] as shown in [13, Fig. 1].

4 1 mmHg ≊ 133.322 Pa.

5 Experimental evidence that the geometry of the in-vivo endfeet configuration matches the geometric assumptions necessary for valve-like motion is still lacking.

6 This does not just hold for our model but models of this kind in general.

